# Unexpected short- and long-term effects of chronic adolescent HU-210 exposure on emotional behavior

**DOI:** 10.1101/2021.12.13.472275

**Authors:** Miguel Farinha-Ferreira, Nádia Rei, João Fonseca-Gomes, Catarina Miranda-Lourenço, Paula Serrão, Sandra H. Vaz, Joana I. Gomes, Valéria Martins, Beatriz de Alves Pereira, Ana M. Sebastião

## Abstract

Chronic adolescent cannabinoid receptor agonist exposure has been shown to lead to persistent increases in depressive-like behaviors. This has been a key obstacle to the development of cannabinoid-based therapeutics. However, most of the published work has been performed with only three compounds, namely Δ9-tetrahydrocannabinol, CP55,940 and WIN55,212-2. Hypothesizing that different compounds may lead to distinct outcomes, we herein used the highly potent CB_1_R/CB_2_R full agonist HU-210, and first aimed at replicating cannabinoid-induced long-lasting effects, by exposing adolescent female Sprague-Dawley rats to increasing doses of HU-210, for 11 days and testing them at adulthood, after a 30-day drug washout. Surprisingly, HU-210 did not significantly impact adult anxious- or depressive-like behaviors. We then tested whether chronic adolescent HU-210 treatment resulted in short-term (24h) alterations in depressive-like behavior. Remarkably, HU-210 treatment simultaneously induced marked antidepressant- and prodepressant-like responses, in the modified forced swim (mFST) and sucrose preference tests (SPT), respectively. Hypothesizing that mFST results were a misleading artifact of HU-210-induced behavioral hyperreactivity to stress, we assessed plasmatic noradrenaline and corticosterone levels, under basal conditions and following an acute swim-stress episode. Notably, we found that while HU-210 did not alter basal noradrenaline or corticosterone levels, it greatly augmented the stress-induced increase in both.

Our results show that, contrary to previously studied cannabinoid receptor agonists, HU-210 does not induce persisting depressive-like alterations, despite inducing marked short-term increases in stress-induced reactivity. By showing that not all cannabinoid receptor agonists may induce long-term negative effects, these results hold significant relevance for the development of cannabinoid-based therapeutics.

## 1. INTRODUCTION

Despite the growing interest in, and therapeutic promise of, endocannabinoid system (ECS) targeting compounds (Black et al., 2019), development of these drugs has been hindered by two factors: for one, activation of the cannabinoid type 1 receptor (CB_1_R) gates not only significant therapeutic, but also psychoactive effects (Solymosi and Kofalvi, 2017). On the other hand, given the well-known role of the ECS in neurodevelopmental processes (Galve-Roperh et al., 2009), there is concern that exposure to these compounds, during such developmental phases, may lead to lasting sequelae (Wong and Wilens, 2017).

Adolescence, a period lasting from years 12-18/25 in humans and post-natal days (PNDs) 28-42/60 in rodents (Schneider, 2013), is one such key developmental window, during which cannabis experimentation also typically begins (Degenhardt et al., 2008). Importantly, the ECS is known to not only play an important role in the neurobiological and behavioral changes that characterize adolescence, but it itself undergoes extensive changes during this period (Meyer et al., 2018). Most notably, CB_1_R density peaks at the onset of adolescence, and gradually decreases towards adulthood, with this developmental trajectory being more evident in the later-maturing prefrontal, and limbic regions possibly accounting for the greater magnitude of CB_1_R-activation mediated effects in the adolescent brain (Rubino and Parolaro, 2011).

Congruently, ample research has focused on the lasting impacts of adolescent cannabis abuse (Higuera-Matas et al., 2015), with findings in humans suggesting negligible long-term impacts on adult anxiety symptomatology (Gobbi et al., 2019), but increased likelihood of depressive and psychotic disorders (Gobbi et al., 2019; Renard et al., 2018), with females being more vulnerable to the deleterious effects on mood (Higuera-Matas et al., 2015; Rubino and Parolaro, 2011). Animal studies have largely corroborated the findings regarding depressive- (Higuera-Matas et al., 2015; Realini et al., 2011) and psychotic-like (Renard et al., 2018) alterations, and the apparent especial vulnerability of females (Higuera-Matas et al., 2015; Rubino and Parolaro, 2011). Furthermore, these changes correlate to significant dysfunctions in glutamatergic (Renard et al., 2017), GABAergic (Renard et al., 2017), dopaminergic (Renard et al., 2017), and endocannabinoid signaling (Lovelace et al., 2015), as well as to marked impairments in synaptic and structural plasticity (Lovelace et al., 2015). As such, and despite the fact that recreational and therapeutic use patterns likely differ (Loflin et al., 2017), information so far available seems to support the notion of adolescence as a period of heightened vulnerability to the deleterious effects of cannabinoids (Meyer et al., 2018; Schneider, 2013), with obvious implications for both recreational and medical users.

However, one limitation of the existing rodent literature on effects of adolescent cannabinoid exposure, relates to the fact that the overwhelming majority of the published work has been performed with only three compounds, namely Δ9-tetrahydrocannabinol (THC; the main psychoactive compound in cannabis (Solymosi and Kofalvi, 2017)), CP55,940, and WIN55,212-2 (Higuera-Matas et al., 2015). Another related limitation is that the results obtained with one of these compounds are often interpreted as holding meaning for other cannabinoid-related drugs. These weaknesses are even more remarkable when considering that not only do different cannabinoid receptor agonists vary in their non-CB_1_R/CB_2_R binding profiles (Pertwee, 2010; Wiley et al., 2016), but also that they are prone to biased agonism, thus triggering differing patterns of CB_1_R-activation-mediated G-protein activation (Diez-Alarcia et al., 2016; Sachdev et al., 2020). As such, the currently available literature is likely an incomplete account of the effects of adolescent cannabinoid exposure. We anticipated that other, yet unstudied, compounds could to lead to distinct outcomes.

Here, we provide a compelling demonstration of this possibility. The work herein reported started as a pilot study where we first aimed at replicating the deleterious effects of adolescent cannabinoid exposure on adult depressive-like behavior in female rats, using the highly potent CB_1_R/CB_2_R full agonist HU-210 (Ferrari et al., 1999; Rodríguez de Fonseca et al., 1996). Unexpectedly, our results showed that, despite having marked short-term effects, chronic adolescent HU-210 exposure did not significantly impact adult anxiety- or depressive-like behavior, nor altered the levels of cannabinoid receptors in two key regions involved in emotional function, the hippocampus and prefrontal cortex (PFC). Furthermore, when comparing outcomes in different tests and interpreting them in light of analytical assays, the work herein reported shows that apparent antidepressant-like effects may be hiding actual prodepressant-like actions. This heavily underlines the necessity of critically assessing the standard interpretations of commonly used behavioral tests, and of using additional sources of data to better interpret results.

## 2. MATERIALS AND METHODS

### 2.1 Subjects and Ethics Approval

Female Sprague-Dawley rats (Charles River Laboratories, Calco, Italy) arrived at the animal facility postnatal day 21 (PND21) and were group housed under standard housing conditions, with *ad libitum* access to food and water, and multiple environmental enrichments. Animals were kept on a 12-hour light-dark cycle (lights on at 6:00AM), with stable temperature (22ºC) and humidity (70%).

All experiments took place during the light phase of the cycle, and were performed in conformity with European Community Guidelines (Directive 2010/63/UE), and with the approval of the Committee for Ethics in Animal Research of the Faculty of Medicine of the University of Lisbon, as well as of the Portuguese Competent Authority for Animal Welfare.

### 2.2. Drugs and Administration

HU-210 [(6aR)-trans-3-(1,1-dimethylheptyl)-6a,7,10,10a-tetrahydro-1-hydroxy-6,6-dimethyl-6H-dibenzo[b,d]pyran-9-methanol; Tocris Bioscience, Bristol, UK] was suspended in dimethyl sulfoxide (DMSO; Sigma-Aldrich, St. Louis, MO, USA) at a 25mM concentration. Aliquots were prepared and stored at −20°C until the day of use, when further dilutions were performed, in 0.9% saline, to reach adequate volume (1ml/kg). At no point was DMSO concentration per injection >1%.

In all experiments, animals were randomly allocated to a treatment condition, and drug administration was performed according to a previously described protocol (Realini et al., 2011): Intraperitoneal (i.p.) HU-210 or vehicle (VEH) injections were administered twice-daily for 11 consecutive days, in an escalating dosing schedule (PND35-37: 25μg/kg; PND38-41: 50μg/kg; PND42-45: 100μg/kg), with 6-7 hours between injections.

### 2.3. Behavioral Testing

Animals were individually handled for at least 5 min/day in the 5 days preceding the start of behavioral testing. On testing days, animals were allowed to acclimatize to the testing room for at least 60 minutes, with testing taking place between 9AM and 6PM. In every test, the testing apparatus was cleaned with 30% ethanol, between animals, to erase olfactory clues.

In the EPM and OFT data was recorded and analyzed using the SMART^®^2.5 video-tracking software (Panlab, Harvard Apparatus, Barcelona, Spain). For the SIT and mFST, videos were recorded and subsequently analyzed, by a trained experimenter blind to drug condition, using Solomon Coder beta 17.03.22 (András Péter, Milan, Italy) behavior coding software.

#### 2.3.1. Elevated Plus Maze (EPM)

The EPM was performed as previously described (Realini et al., 2011). Briefly, animals were individually placed in the center of the EPM apparatus (10 × 50 × 30 cm, raised 50 cm above the floor) facing an open arm, and left to explore the maze for 5 minutes, with the number of entries and time spent in the open arms being recorded.

#### 2.3.2. Open Field Test (OFT)

Testing apparatus consisted of a square black acrylic OF arena (65 × 65 × 40 cm), virtually divided into three concentric square zones. Animals were individually placed into the center of the OF, and allowed to freely explore for 10 minutes. Time spent in the central zone of the OF, as well as total distance travelled and average speed were recorded.

#### 2.3.3. Social Interaction Test (SIT)

The SIT was performed as previously described (Realini et al., 2011). Briefly, after 2 days of habituation to the OF, on the third day, animals were individually placed in the OF with an age-, sex- and weight-matched novel social partner, and were allowed to explore. Time spent, by the test subject, engaging in active social interaction (defined as “sniffing, following, grooming, kicking, mounting, jumping on, wrestling and boxing with, crawling under/over the partner” (File and Seth, 2003)) was recorded over the 10 min trial.

#### 2.3.4. Modified Forced Swim Test (mFST)

The mFST was performed as previously described (Realini et al., 2011). Briefly, a single 15-minute session was performed, during which animals were individually placed into a glass cylinder (20 cm diameter), filled with water at a temperature of 23-25 ºC, to a depth of 30 cm. After each trial, animals were dried with a warm towel, and placed under a heating lamp for at least 10 minutes, and the water was changed.

Time spent engaging in three distinct behaviors was quantified: a) climbing, defined as the rat making “active movements with it forepaws in and out of water” (Detke et al., 1995); b) swimming, defined as the rat making “active swimming motions, more than necessary to merely maintain its head above water” (Detke et al., 1995), which included sporadic bouts of diving and; c) immobility, defined as the rat making “only the movements necessary to keep its head above water” (Detke et al., 1995).

#### 2.3.5. Sucrose Preference Test (SPT)

The SPT was performed as previously described (Realini et al., 2011): animals were individually housed for 72h, and allowed *ad libitum* access to two bottles: one containing tap water, and the other containing a 2% sucrose solution. Bottles were weighed at the start of the test, and every 24 hours afterwards, for the duration of the testing period. To avoid preference or learning effects, the position of the bottles was changed daily, after weighing. For each time-point (24, 48, and 72h) sucrose preference was calculated, being expressed as the percentage of sucrose solution consumed relative to total fluid intake (i.e., values >50% indicate preference for sucrose, and <50% indicate preference for water).

### 2.4. Molecular Analyses

#### 2.4.1. Euthanasia and Sample Collection

At the end of each experiment animals were individually anesthetized with isoflurane, until the paw-pinch reflex was absent, and then decapitated.

Brains were quickly removed, and hippocampus and prefrontal cortex (PFC) samples were isolated in ice-cold artificial cerebral-spinal fluid (aCSF; 124mM NaCl, 3mM KCl, 1.25 mM NaH_2_PO_4_, 26mM NaHCO3, 1mM MgSO_4_, 2mM CaCl_2_, and 10 mM D-glucose, pH 7.4) previously oxygenated with 95% O_2_ and 5% CO_2_. Once isolated, samples were immediately frozen in liquid nitrogen and stored at –80ºC until the moment of use.

Plasma was obtained from trunk blood collected at the moment of sacrifice. For corticosterone analysis, blood was collected into EDTA-coated tubes (Sarstedt AG & Co, Nümbrecht, Germany) and immediately centrifuged at 2000g for 20 minutes at 4 ºC. For noradrenaline analysis, blood was collected into previously heparinized centrifuge tubes, and immediately centrifuged at 1300g for 10 minutes at 4 ºC. In both cases, supernatant was collected after centrifugation and aliquots were stored at –20 ºC until the moment of use.

#### 2.4.2. Western Blotting

Hippocampus and PFC samples were sonicated in Radio-immunoprecipitation Assay (RIPA) buffer containing: 50 mM Tris-HCl (pH 7.5), 150 mM NaCl, 5 mM ethyl-enediamine tetra-acetic acid (EDTA), 0.1% SDS and 1%Triton X-100 and protease inhibitors cocktail (Mini-Complete EDTA-free; Roche Applied Science, Penzberg, Germany). Lysates were centrifuged (13000g, 10min) and the supernatant collected. Supernatant protein concentration was determined through a commercially available Bradford Assay kit (Bio-Rad Laboratories, CA, USA). Equal quantities (50µg) of prepared protein samples were loaded and separated on 10% sodium dodecyl sulphate-polyacrylamide gel electrophoresis (SDS-PAGE) and transferred to polyvinylidene fluoride (PVDF) membrane (GE Healthcare, Buckinghamshire, UK). NZYColour Protein Marker II (NZYTech, Lisbon, Portugal) was used as a protein molecular weight marker. Protein transfer efficacy was confirmed with Ponceau S staining. Membranes were blocked in 3% bovine serum albumin (BSA) in TBS-Tween (20 mM Tris base, 137 mM NaCl and 0.1% Tween-20) for 1h. Membranes were incubated overnight at 4ºC, with guinea pig polyclonal anti-CB_1_R (1:500; CB1-GP-Af530, Frontier Institute Co. Ltd, RRID: AB_2571593) or rabbit polyclonal anti-α-tubulin (1:10000; ab4074, Abcam, RRID: AB_2288001) antibodies diluted in blocking solution, and subsequently incubated with goat polyclonal anti-guinea pig IgG-HRP (1:10000; sc-2438, Santa Cruz Biotechnology RRID: AB_650492) or goat anti-rabbit IgG-HRP (1:10000; #170-6515, Bio-Rad Laboratories; RRID: AB_11125142) secondary antibodies for 1h at RT. Immunoreactivity was detected using Western Lighting ECL Pro (PerkinElmer, MA, USA) and visualized using a Chemidoc XRS+system (Bio-Rad Laboratories, CA, USA). Band intensities were quantified by digital densitometry, using ImageJ 1.52a (National Institutes of Health, Bethesda, MD, USA), with α-tubulin band intensity used as a loading control. Data were normalized for the VEH-treated groups and are expressed as % of vehicle-group protein levels.

#### 2.4.3. ELISA

Plasma corticosterone levels were analyzed using a commercially available Rat Corticosterone ELISA kit (ADI-900-097; Enzo Life Sciences, Farmingdale, NY, USA), according to manufacturer specifications. All samples and standards were run in duplicate, and absorbances read at 405 nm and corrected at 570nm.

#### 2.4.4. HPLC

Plasma noradrenaline levels were assessed through high-performance liquid chromatography with electrochemical detection, as previously described (Soares-da-Silva et al., 1995). Briefly, plasma aliquots were placed in 5ml conical-base glass vials with 50mg alumina and the pH of samples was adjusted to pH 8.3-8.6 by addition of Tris buffer. The adsorbed noradrenaline was then eluted from alumina with 200µl of 0.2M perchloric acid on Costar Spin-X microfilters, 50µl of the eluate was injected into an HPLC (Gilson Medical Electronics, Villiers le Bel, France). The lower limit of detection of noradrenaline was 350fmol.

### 2.5. Bodyweight Changes

The effects of HU-210 exposure on body weight changes were monitored for the duration of the experiments, by subtracting the weight at PND28 to the weight at each subsequent time point, starting with the first day of injections (PND35). To prevent excessive multiple comparisons, the area under the curve (AUC) of the weight changes during drug administration was taken and compared between groups.

### 2.6. Statistical Analysis

Significance level was established at p < 0.05. Outliers were detected using the method outlined by Tukey (Tukey, 1977): observations outside of the interval defined by the first quartile (Q1) - 1.5 interquartile range (IQR) and the third quartile (Q3) + 1.5 IQR, were considered outliers and removed from analysis. Data was analyzed through two-tailed unpaired Student’s *t*-tests, or Two-Way Analysis of Variance (ANOVA), with Holm-Sidak correction for multiple comparisons when appropriate, and are expressed as mean ± standard error of mean (SEM). All statistical analyses were performed using GraphPad Prism 8 software (GraphPad Software, San Diego, CA, USA).

## 3. RESULTS

### 3.1. Experiment 1. Chronic adolescent HU-210 exposure does not induce long-lasting behavioral or molecular changes

To determine whether chronic adolescent (PND35-45) HU-210 exposure led to significant alterations in adult anxiety- and depression-like behaviors, animals were tested after a 30-day washout period (PND76; Fig. 1A), in the elevated plus maze (EPM), open field (OFT), social interaction (SIT), modified forced swim (mFST), and sucrose preference tests (SPT).

**Figure 1.**
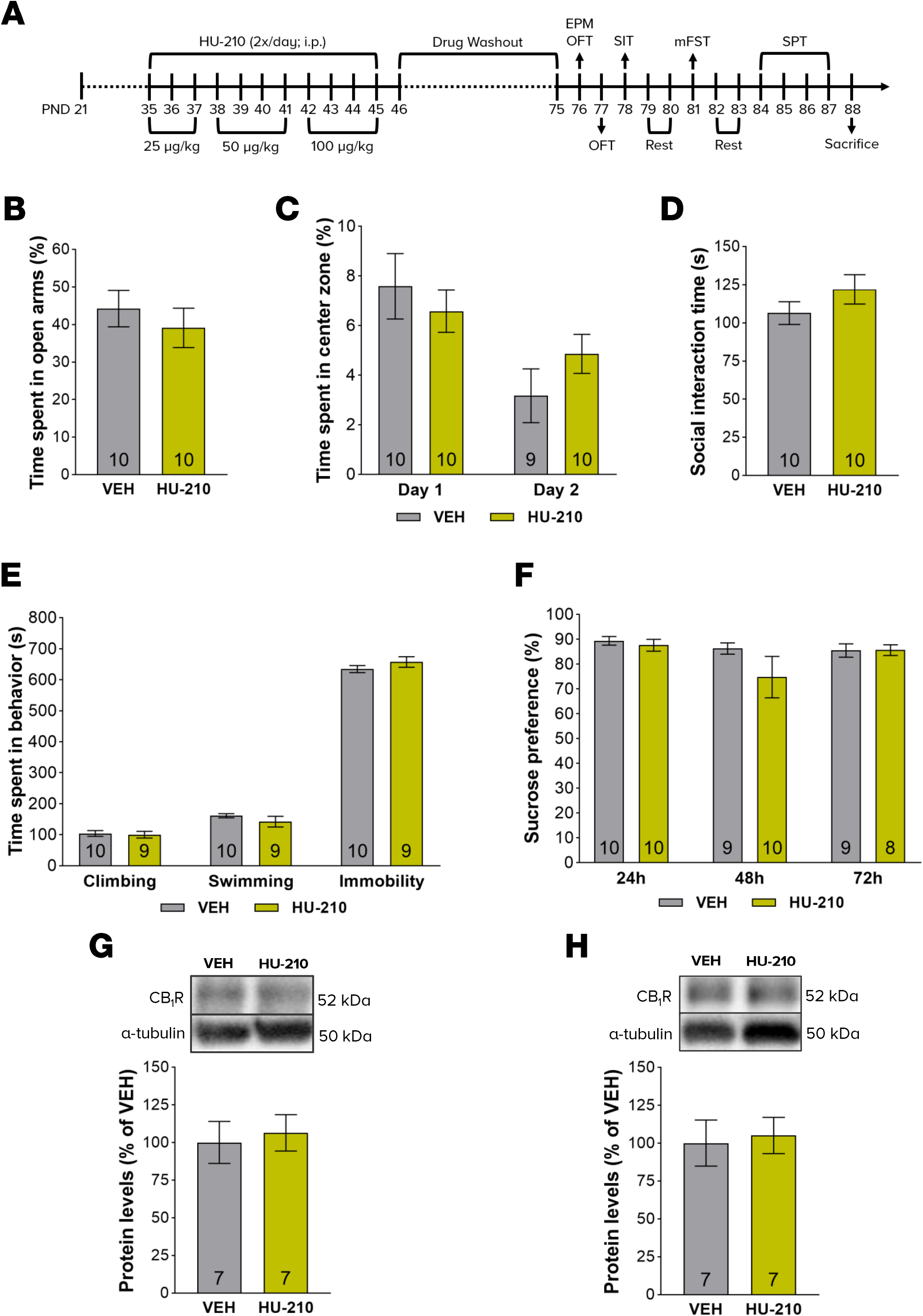
Chronic adolescent HU-210 exposure does not alter adult anxiety- or depression-like behaviors, nor hippocampal or prefrontal CB_1_R protein levels. **A**. Timeline of experimental procedures. Adolescent female Sprague-Dawley rats received twice-daily intraperitoneal injections of ascending doses of HU-210 (PND35-37: 25µg/kg, PND38-41: 50µg/kg, PND42-45: 100µg/kg) or equivalent vehicle, for a period of 11 days. After a 30-day drug washout, animals were tested in a battery of behavioral tests of anxiety- (EPM, OFT, SIT) and depression-like (FST, SPT) behaviors. Following the end of testing, animals were sacrificed and hippocampal and prefrontal cortex tissue samples were collected for Western Blot; **B-D**. Adolescent HU-210 treatment did not induce long-lasting alterations in anxiety-like behavior, as shown by the absence of differences between groups in percentage of time spent in the open arms of the EPM **(B)**, percentage of time spent in the center zone of the open field **(C)**, and time spent in social interaction in the SIT **(D)**; **E-F**. Animals chronically exposed to HU-210 as adolescents performed similar to vehicle-treated controls in all three mFST parameters **(E)**, and all three SPT time-points **(F)**, suggesting no long-term treatment effects upon depressive-like behaviors. **G-H**. No treatment-induced differences were observed in either adult hippocampal or prefrontal CB_1_R protein levels. Data are expressed as mean ± SEM, with number of animals *n* indicated at the bottom of plot bars; unpaired student’s t-test or repeated measures Two-way ANOVA, with Holm-Sidak correction for multiple comparisons where appropriate.

In the EPM, HU-210-exposed rats did not present altered anxiety-like behavior, with no difference being observed between groups regarding either percentage of time spent [t(18) = 0.72, p = 0.483; Fig. 1B], or number of entries [t(14) = 0.85, p = 0.41; Fig. S1B], in the open arms. Accordingly, no statistical difference was observed in center zone permanence in either the first [t(18) = 0.642, p = 0.529] or second [t(17) = 1.28, p = 0.217] OFT sessions (Fig. 1C), nor in time spent engaging in active social interaction in the SIT [t(18) = 1.3, p = 0.217; Fig. 1D]. Importantly, the absence of changes was not attributable to alteration in locomotor activity, insofar no differences were observed in total distance travelled in either the first [t(18) = 0.91, p = 0.144] or second [t(18) = 0.027, p = 0.979] OFT sessions (Fig. S1C).

Surprisingly, HU-210 treatment had no significant effect upon mFST performance (Fig. 1E), with HU-210 and vehicle-treated animals spending similar time climbing [t(17) = 0.266, p = 0.793], swimming [t(17) = 1.08, p = 0.297], and immobile [t(17) = 1.14, p = 0.27]. Furthermore, repeated measures two-way ANOVA found no statistically significant effect of HU-210 treatment on sucrose preference [F(1, 18) = 1.27; p = 0.275; Fig. 1F].

At the end of behavioral testing, animals were sacrificed, and hippocampal and PFC samples were collected for western blot analysis of CB_1_R protein levels. Congruently with behavioral data, adolescent HU-210 treatment did not significantly alter adult hippocampal [t(12) = 0.35, p = 0.734; Fig. 1G] or prefrontal [t(12) = 0.26, p = 0.798; Fig. 1H] CB_1_R protein levels.

### 3.2. Experiment 2. Chronic adolescent HU-210 exposure induces short-term antidepressant-like effects in the mFST

The lack of long-term effects observed in experiment 1 was somewhat unexpected, given the previous literature with other cannabinoid-related compounds (Higuera-Matas et al., 2015; Realini et al., 2011). That HU-210 treatment was biologically active, could however be concluded from the fact that it induced a significant decrease in weight gain [t(198) = 7.5, p = 0.001; Fig. S1A]. We thus aimed at determining whether the absence of HU-210 effects in experiment 1 was due to the compound not affecting behavior, or to the washing out of effects during the 30-day rest period. For this we repeated the drug administration protocol with another set of animals, but performed the OFT and mFST 24h after the last injection (PND46; Fig. 2A).

**Figure 2.**
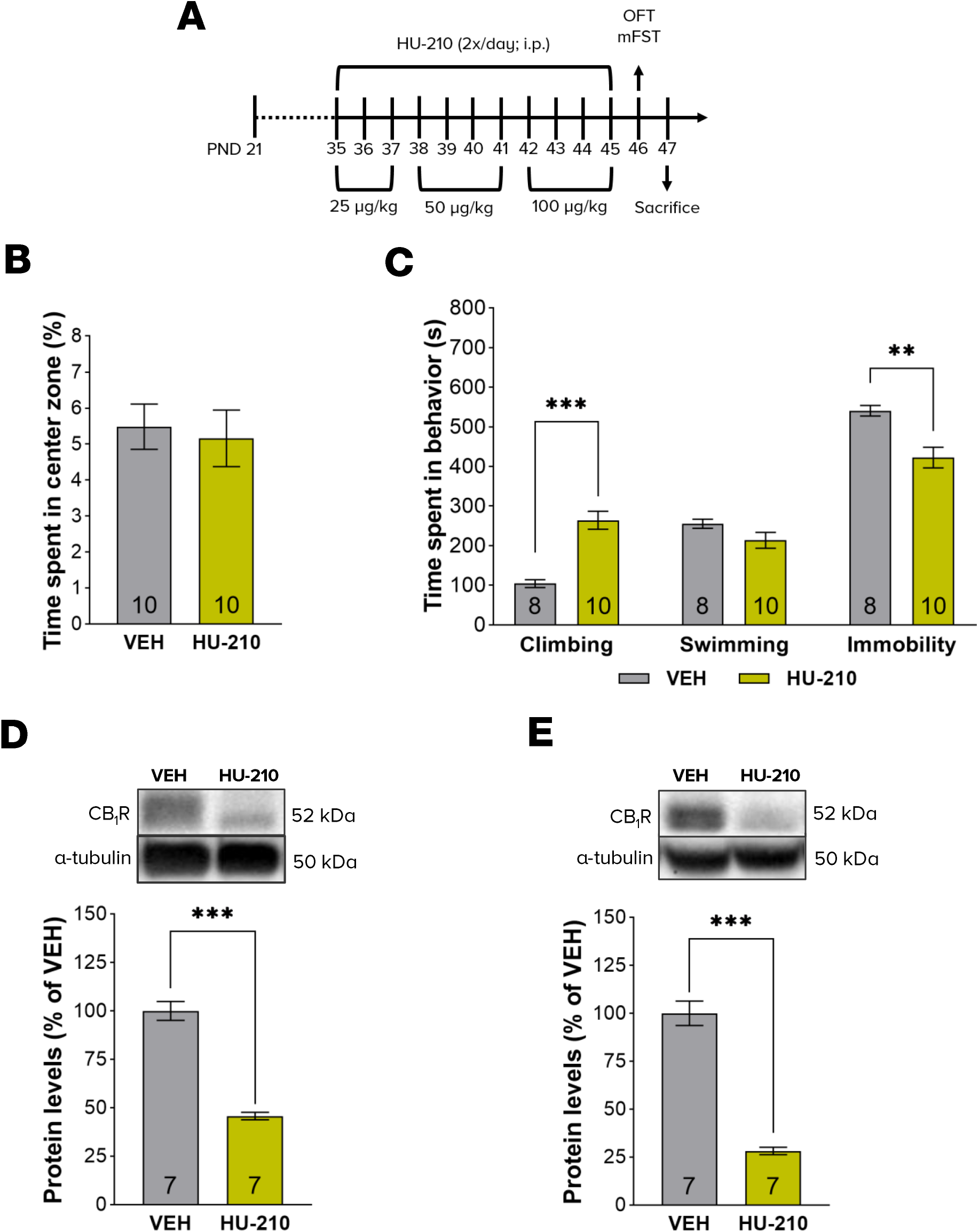
Chronic adolescent HU-210 exposure induces significant short-term decreases in depression-like behavior and hippocampal and prefrontal CB_1_R protein levels, without affecting anxiety-like behavior. **A**. Timeline of experimental procedures. Adolescent female Sprague-Dawley rats received twice-daily intraperitoneal injections of ascending doses of HU-210 (PND35-37: 25µg/kg, PND38-41: 50µg/kg, PND42-45: 100µg/kg) or equivalent vehicle, for a period of 11 days. 24-hours after the last injection, animals were tested in the OFT and mFST, respectively assessing anxiety- and depression-like behaviors. On the day following behavioral testing (PND47) animals were sacrificed, and tissue samples were collected from the hippocampus and PFC for Western Blot; **B**. Adolescent HU-210 treatment did not alter the percentage of total time spent in the center zone of the open field, thereby suggesting treatment did not alter anxiety-like behavior; **C**. Adolescent HU-210 treatment had significant antidepressant-like effects in the mFST, as shown by significantly increased climbing time, and concomitantly decreased immobility time, without altering time spent swimming; **D-E**. Hippocampal and prefrontal CB_1_R protein levels were markedly reduced by chronic adolescent HU-210 exposure. Data are expressed as mean ± SEM, with number of animals *n* indicated at the bottom of plot bars; ** p ≤ 0.01, *** p ≤ 0.001, unpaired student’s t-test.

Remarkably, in the mFST (Fig. 2C), HU-210-treated animals evidenced a marked increase in climbing time [t(16) = 5.95, p < 0.001], and a concomitantly marked decrease in immobility [t(16) = 3.74, p = 0.002], with no alterations in swimming [t(16) = 1.71, p = 0.107], suggesting an antidepressant-like effect. Furthermore, this effect was not attributable to alterations in either anxiety-like behavior or spontaneous locomotor activity, as neither OFT center zone permanence [t(18) = 0.32, p = 0.753; Fig. 2B], nor total distance travelled [t(17) = 2, p = 0.059; Fig. S2B] were found to be significantly changed.

On the day following behavioral testing (PND47), animals were sacrificed, and hippocampus and PFC samples were collected for western blot. These analyses revealed a marked HU-210-induced decrease in both hippocampal [t(12) = 10, p = 0.001; Fig. 2D] and prefrontal [t(12) = 11, p < 0.001; Fig. 2E] CB_1_R protein levels.

### 3.3. Experiment 3. Chronic adolescent HU-210 exposure induces short-term prodepressant-like effects in the SPT

Results in experiment 2 suggested HU-210 treatment induced a short-term antidepressant-like effect. However, despite the lack of changes in locomotor activity (Fig. S2B), personal observation and manipulation of the animals suggested that HU-210-treated rats were markedly hyperreactive (Ferrari et al., 1999; Rodríguez de Fonseca et al., 1996), suggesting a possible biasing influence to mFST results (Ferreira et al., 2018). As such, to determine whether the results of experiment 2 represented an actual antidepressant-like effect, a new group of animals was manipulated as before, and tested in the SPT – a test of depressive-like behavior not relying on locomotor function – and the EPM. As in experiment 2, testing started 24h after the last HU-210 administration (PND46-49; figure 3A).

**Figure 3.**
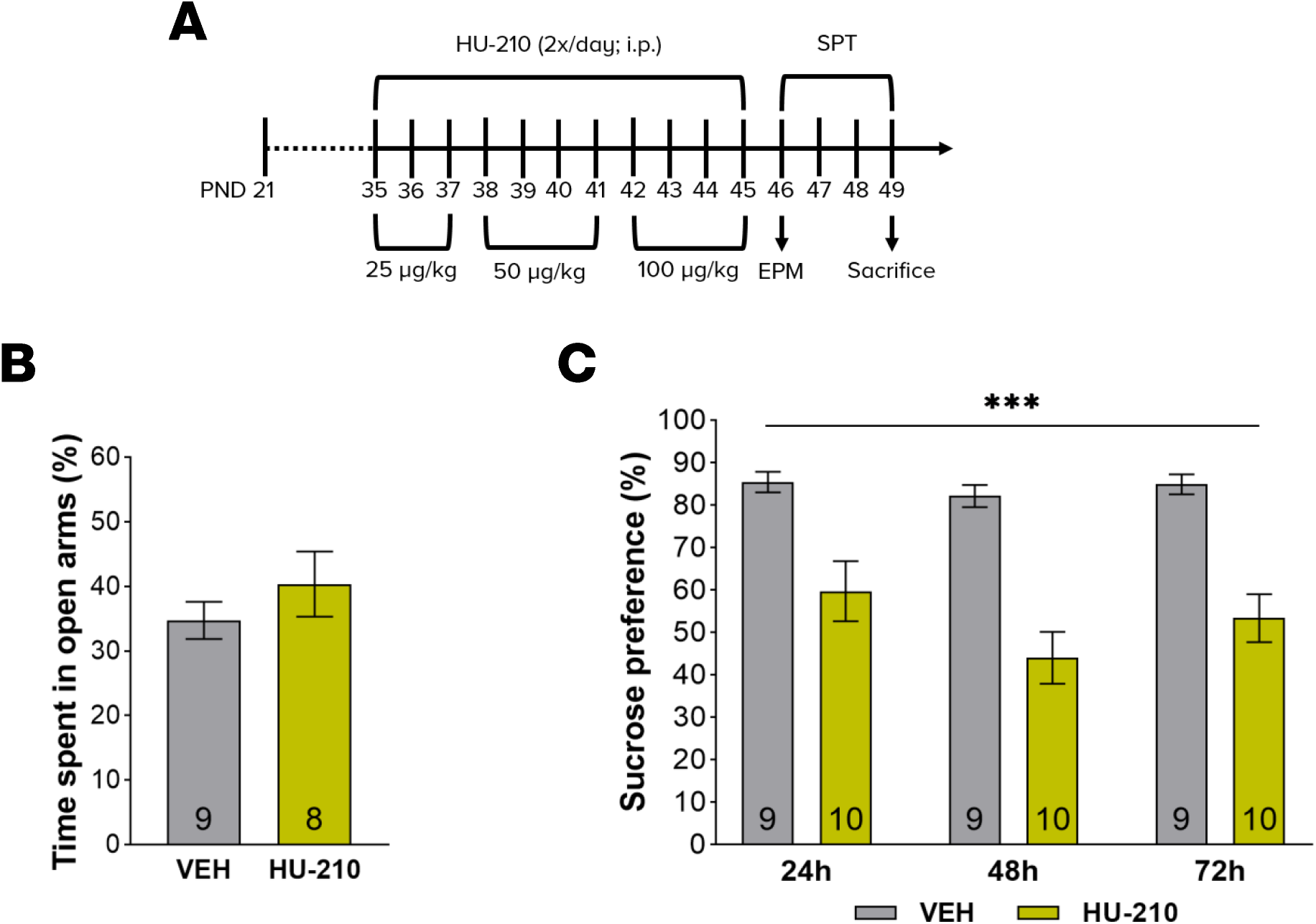
Chronic adolescent HU-210 exposure induces significant short-term increases in depression-like, but not anxiety-like, behaviors. **A**. Timeline of experimental procedures. Adolescent female Sprague-Dawley rats received twice-daily intraperitoneal injections of ascending doses of HU-210 (PND35-37: 25µg/kg, PND38-41: 50µg/kg, PND42-45: 100µg/kg) or equivalent vehicle, for a period of 11 days. Starting 24-hours after the last injection, animals were tested in the EPM and SPT, respectively assessing anxiety- and depression-like behaviors; **B**. Adolescent HU-210 treatment did not alter the percentage of total time spent in the open arms of the EPM, suggesting treatment did not alter anxiety-like behavior; **C**. HU-210-treated animals showed marked and consistent decreases in sucrose preference across the three time-points, suggesting a pronounced prodepressant-like effect of HU-210 treatment. Data are expressed as mean ± SEM, with number of animals *n* indicated at the bottom of plot bars; *** p ≤ 0.001, unpaired student’s t-test or repeated measures Two-way ANOVA, with Holm-Sidak correction for multiple comparisons where appropriate.

Here, in contrast with the results of experiment 2, repeated measures two-way ANOVA found a significant effect of HU-210 treatment on sucrose preference [F(1, 17) = 29.1; p = 0.001; Fig. 3C], with HU-210 animals showing significantly decreased sucrose preference, relative to their vehicle-treated counterparts, suggesting a prodepressant-like effect. In the EPM, no significant differences were detected in either the percentage of time spent [t(15) = 1, p = 0.335; Fig. 3B], or the number of entries [t(16) = 1.3, p = 0.219; Fig. S3B], in the open arms.

### 3.4. Experiment 4. Chronic adolescent HU-210 exposure induces contradictory effects in the mFST and SPT

Given the contradictory nature of the results of experiments 2 and 3, we wanted to control for the possibility that the discrepancy resulted from variation between different sets of animals. To this end, a new group of animals was treated as previously described, and then tested in both the mFST and the SPT starting 24h after the last HU-210 injection (PND46-49; figure 4A)

**Figure 4.**
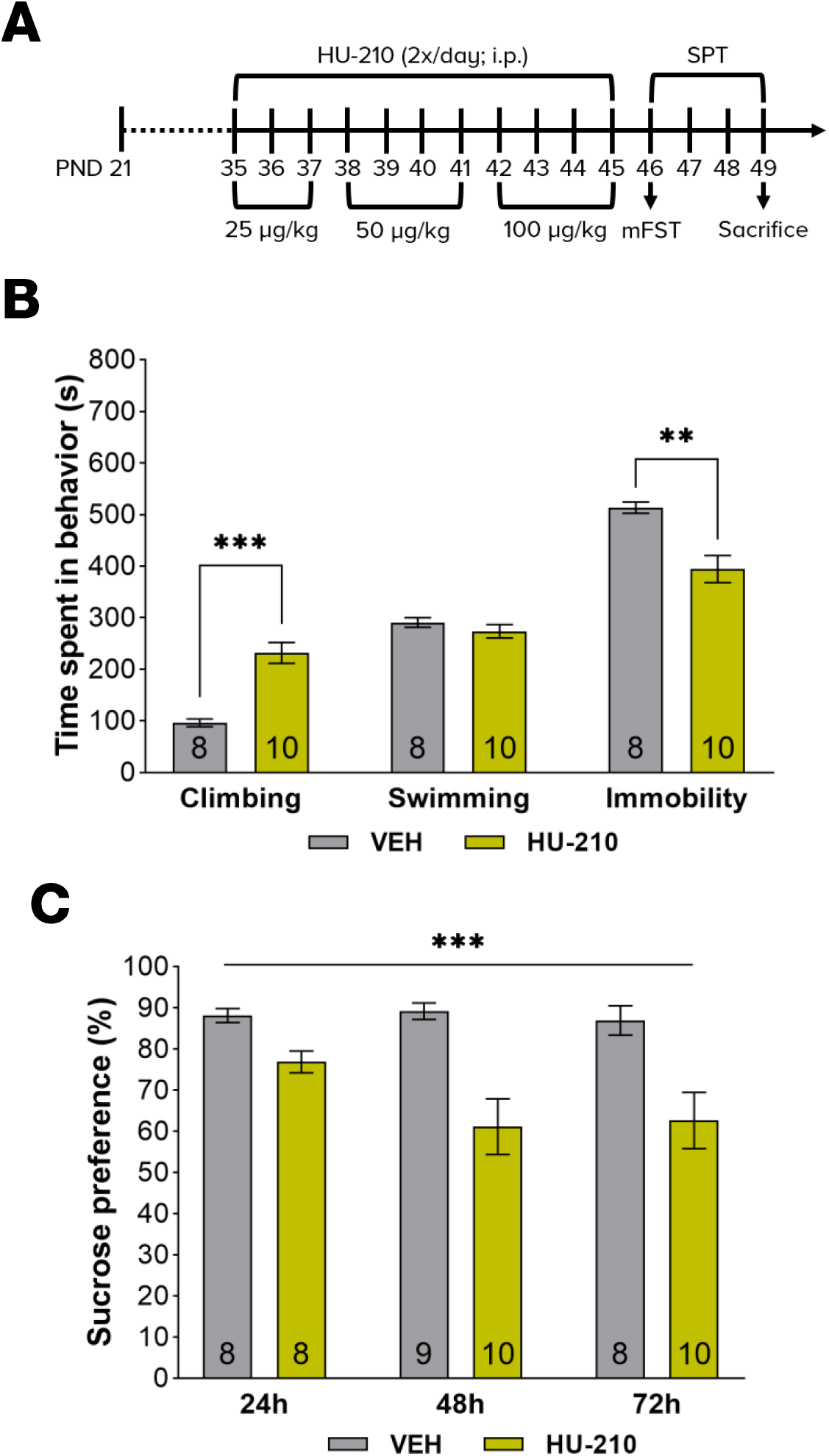
Chronic adolescent HU-210 exposure induces contradictory short-term alterations in depression-like behaviors. **A**. Timeline of experimental procedures. Adolescent female Sprague-Dawley rats received twice-daily intraperitoneal injections of ascending doses of HU-210 (PND35-37: 25µg/kg, PND38-41: 50µg/kg, PND42-45: 100µg/kg) or equivalent vehicle, for a period of 11 days. Starting 24-hours after the last injection, the same batch of animals were tested in both the mFST and SPT; **B**. Adolescent HU-210 treatment induced significant increases in time spent climbing, and concomitant decreases in immobility time, without altering time spent swimming, suggesting a marked antidepressant-like effect; **C**. On the other hand, in the SPT, the HU-210-treated group evidenced a marked decrease in sucrose preference, across all time points, suggesting a prodepressant-like action of HU-210 treatment. Data are expressed as mean ± SEM, with number of animals n indicated at the bottom of plot bars; ** p ≤ 0.01, *** p ≤ 0.001, unpaired student’s t-test or repeated measures Two-way ANOVA, with Holm-Sidak correction for multiple comparisons where appropriate

In the mFST (Fig. 4B), we replicated the findings of experiment 2, whereby HU-210 treatment induced a marked increase in climbing [t(16) = 5.64, p < 0.001], and decrease in immobility [t(16) = 3.81, p = 0.002], without altering the time spent swimming [t(16) = 1, p = 0.331], suggesting an antidepressant-like effect. Interestingly, and despite this, we also replicated the findings of experiment 3, as HU-210-treated animals showed significantly diminished preference for sucrose in the sucrose preference test [F(1, 17) = 15.3; p = 0.001; Fig. 4C], suggesting a prodepressant-like effect.

### 3.5. Experiment 5. Chronic adolescent HU-210 exposure increases the amplitude of the neurochemical response to acute stress

Knowing that the discrepancy in the outcomes from the mFST and SPT was not an artifact of animal variation, we aimed at understanding their neurochemical underpinnings. To this end, a final group of animals was manipulated as before, and 24h after the last injection we sacrificed half of the animals treated with vehicle- or HU-210, with blood being rapidly collected to assess basal levels of plasma corticosterone and noradrenaline, via ELISA and HPLC, respectively. The remaining animals in each treatment group were sacrificed 30 minutes (Connor et al., 1997) after being exposed to a 15-minute swim stress, replicating mFST conditions, to assess stress-induced changes in plasma corticosterone and noradrenaline levels.

Two-way ANOVA of plasma corticosterone levels (Fig. 5B), revealed significant main effects of both treatment [F(1, 19) = 8.68; p = 0.008] and stress [F(1, 19) = 22.1; p < 0.001]. Additionally, a significant treatment x stress interaction was observed [F(1, 19) = 5.75; p = 0.027], suggesting the effects of swim stress differ as a function of treatment. Indeed, whereas swim stress did not lead to a significant increase in plasma corticosterone levels in vehicle-treated animals [t(19) = 1.59; p = 0.337], it did so in HU-210-treated ones [t(19) = 5.14; p = < 0.001]. Furthermore, stress-exposed HU-210 animals evidenced higher plasma corticosterone levels than their vehicle-treated counterparts [t(19) = 3.69; p = 0.006], despite there being no differences in basal corticosterone levels between treatment groups [t(19) = 0.397; p = 0.696].

**Figure 5.**
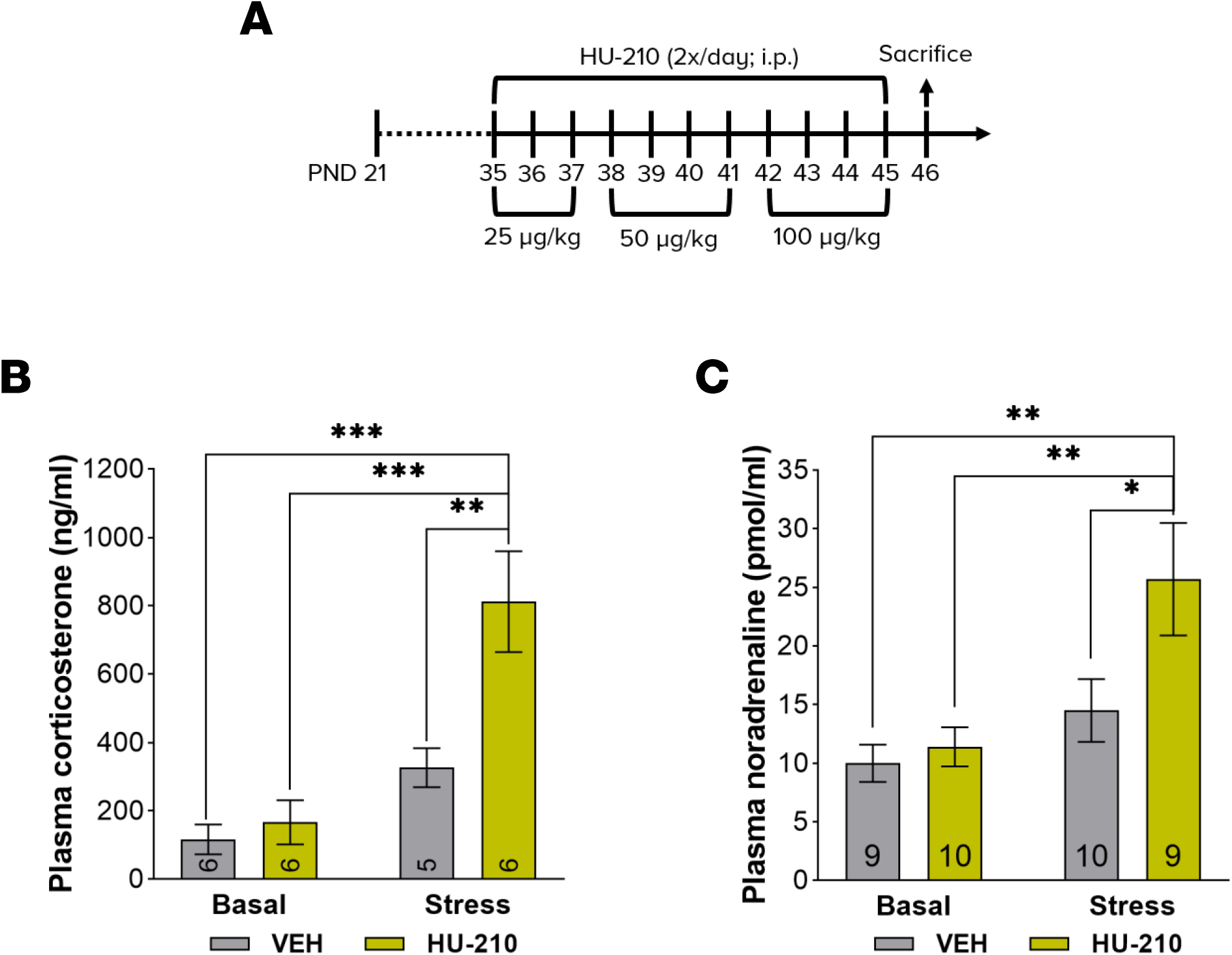
Chronic adolescent HU-210 exposure magnifies the effects of acute stress on noradrenaline and corticosterone levels, without altering basal levels. **A**. Timeline of experimental procedures. Adolescent female Sprague-Dawley rats received twice-daily intraperitoneal injections of ascending doses of HU-210 (PND35-37: 25µg/kg, PND38-41: 50µg/kg, PND42-45: 100µg/kg) or equivalent vehicle, for a period of 11 days. 24-hours after the last drug injection, animals were either sacrificed immediately, to assess basal plasmatic levels of noradrenaline and corticosterone, or 30-minutes after a 15-min swim stress exposure to assess stress-induced changes in plasmatic noradrenaline and corticosterone levels; **B**. No differences were observed between treatment groups in basal plasma corticosterone levels. However, swim stress exposure led to a marked increase in plasma corticosterone levels in HU-210-treated animals, whereas in vehicle treated controls no such effect was observed; **C**. Similarly, while no treatment-induced differences were observed in basal plasma noradrenaline levels, a significant difference emerged after swim-stress exposure, whereby the HU-210, but not the vehicle-treated, group evidenced a significant increase in plasma noradrenaline levels. Data are expressed as mean ± SEM, with number of animals n indicated at the bottom of plot bars; * p < 0.05, ** p ≤ 0.01, *** p ≤ 0.001, Two-way ANOVA with Holm-Sidak correction for multiple comparisons.

Moreover, two-way ANOVA of plasma noradrenaline (Fig. 5C) found significant main effects for both treatment [F(1, 34) = 4.69; p = 0.037] and stress [F(1, 34) = 10.4; p = 0.003], though the interaction was not significant [F(1, 34) = 2.84; p = 0.101]. Post-hoc comparisons revealed that while swim stress significantly increased plasma noradrenaline levels in HU-210-treated animals [t(34) = 3.47; p = 0.007], it did not significantly do so in vehicle-treated ones [t(34) = 1.09; p = 0.631]. Furthermore, comparison of stress-exposed vehicle- and HU-210-treated groups showed that HU-210 treated animals exposed to the swimming stress had significantly higher plasma noradrenaline levels [t(34) = 2.72; p = 0.04], than stress-exposed vehicle-treated animals. Contrastingly, in animals not exposed to the swimming stress, HU-210 treatment did not significantly increase plasma noradrenaline [t(34) = 0.34; p = 0.736].

## 4. DISCUSSION

The main findings of the present work are that while chronic adolescent HU-210 exposure did not induce long-term changes in behavioral tests of depressive-like behavior, it induced marked short-term anti- and prodepressant-like effects in the mFST and SPT, respectively. Importantly, HU-210 treatment also clearly increased short-term stress reactivity, which likely biased mFST results.

Adolescent exposure to cannabinoid receptor agonists has been consistently found to induce long-lasting deleterious effects upon adult affective behavior (Higuera-Matas et al., 2015; Realini et al., 2011). While aiming at replicating this effect, using the CB_1_R/CB_2_R full-agonist HU-210, we unexpectedly found that chronic adolescent HU-210 exposure had no significant impact on adult anxiety- or depression-like behaviors, nor did it alter hippocampal or prefrontal CB_1_R protein levels, in female rats. Such findings, while boding well for the development of cannabinoid-based therapies, are nonetheless in stark contrast with the literature on adolescent cannabinoid exposure (Higuera-Matas et al., 2015; Realini et al., 2011). In fact, the only previous study of adolescent HU-210 exposure (Lee et al., 2014), reports an increase in stress reactivity of adult male rats exposed to acute restraint stress, whereas in females an impairment was observed in adult hippocampal neurogenesis. Both findings are consistent with prodepressant-like actions of HU-210. On the other hand, in studies with adult HU-210 exposure, significant antidepressant-, rather than prodepressant-like, effects were reported (Jiang, 2005; Morrish et al., 2009).

If relying only in the data obtained with the mFST 24h after HU-210 administration, we would conclude that adolescent HU-210 exposure induces short-term antidepressant-, rather than prodepressant-like effects. This conclusion, while surprising, would be consistent with the previously mentioned studies of adult HU-210 exposure (Jiang, 2005; Morrish et al., 2009). However, and despite no obvious differences in either anxiety-like behavior or locomotor activity indexes being observed, HU-210-treated animals evidenced markedly hyperreactive behavior, which could confound mFST performance (Ferreira et al., 2018). Notably, such hyperreactivity had already been described with adult HU-210-exposed animals, as shown by increased plasma corticosterone levels, as well as altered behavioral performance and vocalizations in response to acute stress (Ferrari et al., 1999; Hill and Gorzalka, 2006; McLaughlin et al., 2009; Rodríguez de Fonseca et al., 1996). Importantly, none of these studies assessed plasma noradrenaline levels (Ferrari et al., 1999; Hill and Gorzalka, 2006; McLaughlin et al., 2009; Rodríguez de Fonseca et al., 1996). Behavioral reactivity is unlikely to affect performance in the SPT (Ferreira et al., 2018), and data from this test clearly showed that HU-210 causes a marked, albeit non-persistent, prodepressant-like effect. Importantly, the apparent contradicting results in the SPT and mFST, could be found while testing different batches of animals, as well as within the same group of animals. Thus, assuming that the same treatment cannot simultaneously induce antidepressant- and prodepressant-like effects, the data suggested that one of the two results was a false-positive.

In light of the increased hyperreactivity to stress caused by HU-210, the false-positive result is likely the one derived from the mFST. Specifically, increased climbing in the mFST has been associated with an increase in noradrenergic signaling (Detke et al., 1995), and HU-210-induced increases in climbing have been shown to be respectively attenuated or abolished by α_1_ and β receptor antagonists (Morrish et al., 2009). Thus, we hypothesized that HU-210 treatment would lead to abnormally high noradrenaline levels, possibly due to an impairment in the enzymatic inactivation of noradrenaline (Abboussi et al., 2020), which in conjunction with increased sensitivity of noradrenergic receptors (Reyes et al., 2012), would result in exaggerated behavioral reactivity, leading to increased climbing in the mFST. In fact, in spite of an increase in noradrenergic signaling being associated with antidepressant-like responses (Cryan et al., 2005), it must be noted that increased noradrenaline levels have been observed in the CSF of depressed patients (Potter and Manji, 1994), and that increased noradrenergic signaling was implicated in mediating the aversive properties of cannabinoid receptor agonists (Carvalho et al., 2010; Carvalho and Van Bockstaele, 2011). Furthermore, not only does the ECS directly modulate hypothalamic-pituitary-adrenal (HPA) axis activity (Martın-Calderon et al., 1998; Micale and Drago, 2018), but there also is a reciprocal relationship between noradrenergic and HPA axis functioning (Dunn and Swiergiel, 2008). Noradrenaline receptor antagonists have also been shown to blunt HU-210-induced hyperreactivity to stress (McLaughlin et al., 2009). Remarkably, and in accordance with our hypothesis that the outcome from the mFST was a false positive, while adolescent HU-210-exposuret did not significantly increase plasmatic levels of either corticosterone or noradrenaline under basal conditions, it greatly magnified the amplitude of the stress-induced increase in the levels of both. As such, our results clearly demonstrate that HU-210 treatment selectively increased stress-reactivity in adolescent rats, while simultaneously identifying the neurochemical correlates of such finding. Since the mFST triggers an exaggerated behavioral climbing response, it can be mistakenly identified as an antidepressant-like effect. Interestingly, our data is also consistent with a recent suggestion that the mFST may be better interpreted as a test of anxiety-like behavior, than of depressive-like behavior (Anyan and Amir, 2018).

This misleading mFST performance is in line with that previously observed with the CB_1_R antagonist/inverse agonist rimonabant (Griebel et al., 2005), now known to have prodepressant effects (Boekholdt and Peters, 2010). This, combined with our findings that both hippocampal and prefrontal CB_1_R protein levels were markedly reduced shortly after HU-210 exposure, suggests that the HU-210-induced prodepressant-like effects may, at least partially, derive from HU-210-induced CB_1_R downregulation, thus functionally similar to receptor antagonism. Indeed, the ECS is known to act in the hippocampus and PFC as a brake to the stress response (Morena et al., 2016). Furthermore, insofar both the PFC (Renard et al., 2017) and the hippocampus (Floresco et al., 2001) exert regulatory actions over the mesolimbic reward circuitry, such a disruption of normal CB_1_R-mediated signaling in both regions, might account for the anhedonic-like effects observed in the SPT. Importantly, it must be noted that it is highly unlikely that the observed effects are due to the effects of acute withdrawal, insofar similar work with adult rats chronically exposed to HU-210, found that HU-210-maintained animals retained the increased climbing in the mFST (Morrish et al., 2009).

Besides the mesolimbic, prefrontal and hippocampal areas, other brain regions are likely involved in the short-lasting prodepressant-like action of HU-210 exposure. For example, previous work from our lab has shown that adult mice chronically exposed to WIN55,212-2 present alterations in lateral habenula (LHb) metabolism and connectivity (Mouro et al., 2018). Congruently, CB_1_R downregulation-induced changes in LHb excitability, known to be associated with depressive-like effects (Berger et al., 2018; Park et al., 2017), might contribute to decreased sucrose preference. Moreover, cannabinoid receptor agonist administration is known to have significant aversive properties (Carvalho et al., 2010; Carvalho and Van Bockstaele, 2011). It is therefore plausible that chronic HU-210 administration may, in and of itself, work as a chronic stressor, which could contribute to the short-term effects in the SPT. Indeed, such an assertion is further supported by the fact that HU-210 administration reliably induced a reduction of weight gain across different cohorts of animals (see Supplementary Figs. S1A, S2A, S3A & S4A). While this hypothesis could conceivably be tested by co-administering HU-210 with a CB_1_R antagonist, in reality it is likely to not be as simple, given the known deleterious impact of CB_1_R antagonists on mood (Beyer et al., 2010; Boekholdt and Peters, 2010).

An issue that still deserves discussion is the lack of long-term effects of HU-210, vis a vis other, less potent/effective, cannabinoid receptor agonists. Indeed, this is even more surprising when considering that we used female animals, which are generally reported as being most vulnerable to such long-term affective behavioral effects (Higuera-Matas et al., 2015; Rubino and Parolaro, 2011). The most compelling possibility is that the difference in the long-term effects may result from subtle distinctions in the pharmacodynamics of HU-210, be they in its binding to non-ECS receptors (Pertwee, 2010; Wiley et al., 2016), or in the signaling pathways triggered by CB_1_R binding (Diez-Alarcia et al., 2016; Sachdev et al., 2020).

Independently of the causes, the variability in the long-lasting effects of cannabinoid compounds reinforces the need to diversify the pool of drugs used for ECS research, to allow disclosure of the full range of possible effects of ECS manipulation. This has significant impact on the development of cannabinoid-based therapeutics, and when addressing the sequelae resulting from the abuse of an increasingly diverse range of synthetic cannabinoid compounds.

Another important conclusion of this work also pertains to the need for careful interpretation of data derived from behavioral methodologies. Our results call attention to the need to carefully consider results obtained with any given behavioral test, in light not just of those results obtained with other tests, but also in light of the behavior of the animals in non-test environments. Indeed, our observation that HU-210 animals behaved in a markedly altered fashion during everyday manipulation led us to a critical interpretation of the mFST results, and to directly address the possibility of enhanced reactivity to stress, by searching for the subjacent neurochemical correlates. Thus, our results cement the usefulness of biological data as tools to better refine and narrow the range of possible interpretations for any given behavioral output.

In conclusion, we have shown that chronic adolescent exposure to a widely used high affinity CB_1_R/CB_2_R agonist induces a markedly different, and complex, pattern of behavioral and neurobiological alterations, which hold significant relevance for the development of cannabinoid-based therapeutics, and for the study of the ECS under both physiological and pathological situations.

## 6. AUTHOR CONTRIBUTIONS

MF-F and AMS conceived and designed the experiments. MF-F, performed and analyzed the behavioral experiments. VM and BP provided assistance during behavioral experiments and animal manipulation. MF-F, NR, SHV, and JIG, performed the dissections for sample acquisition. NR, JF-G and CM-L, performed the western blots. NR performed the ELISA. PS performed the HPLC assays. MF-F and AMS performed the statistical analyses, and wrote the manuscript in consultation with the remaining authors.

## 7. FUNDING & ACKNOWLEDGEMENTS

This work was supported by project funding from Fundação para a Ciência e para a Tecnologia (FCT) to AMS (PTDC/MED-FAR/30933/2017). This project has received funding from H2020-WIDESPREAD-05-2017-Twinning (EpiEpinet) under grant agreement No. 952455. MF-F (SFRH/BD/147505/2019), NR (PD/BD/113463/2015), JF-G (PD/BD/114441/2016) and CM-L (SFRH/BD/118238/2016) are supported by PhD fellowships from FCT.

The authors would like to thank Dr. Attila Köfalvi, PhD at the Center for Neuroscience and Cell Biology of Coimbra, Faculty of Medicine, University of Coimbra (Coimbra, Portugal) for the insightful informal discussions.

