## Supplemental figures for "Unexpected short- and long-term effects of chronic adolescent HU-210 exposure on emotional behavior"

### SUPPLEMENTARY FIGURES

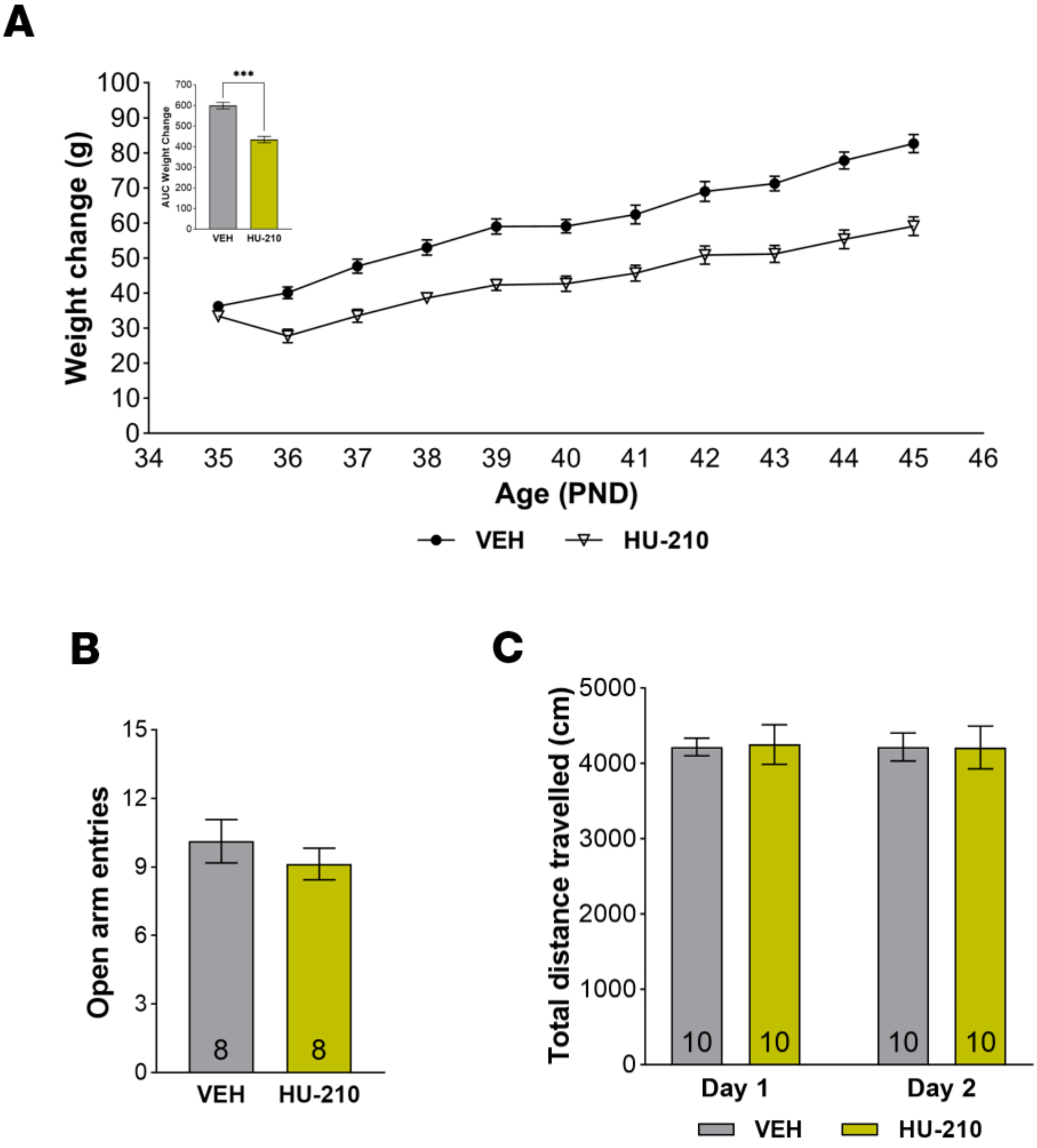


**Supplementary Figure S1. Chronic adolescent HU-210 treatment induces significant decreases in weight-gain, without affecting adult anxiety-like behavior or spontaneous locomotor activity.**

**A.** Changes in animal weight over time. To assess alterations in weight gain, the weight of each animal at PND28 was subtracted to its daily weight during the period of injections (PND35-45). To avoid multiple comparisons, the area under curve was obtained and compared between groups (inset), and evidenced a clear HU-210-induced decrease in weight-gain; **B.** In accordance with the absence of differences in open arm permanence in the EPM (see fig. 1B), no differences were observed in the number of open arm entries; **C.** HU-210-treated animals did not significantly differ from vehicle-treated controls with regards to the total distance travelled during either the first or the second OFT trial, suggesting treatment did not induce any lasting changes in locomotor activity. Data are expressed as mean ± SEM, with number of animals *n* indicated at the bottom of plot bars; *** $p \leq 0.001$, unpaired student's t-test. Graphs correspond to the animals in experiment 1 (see text).

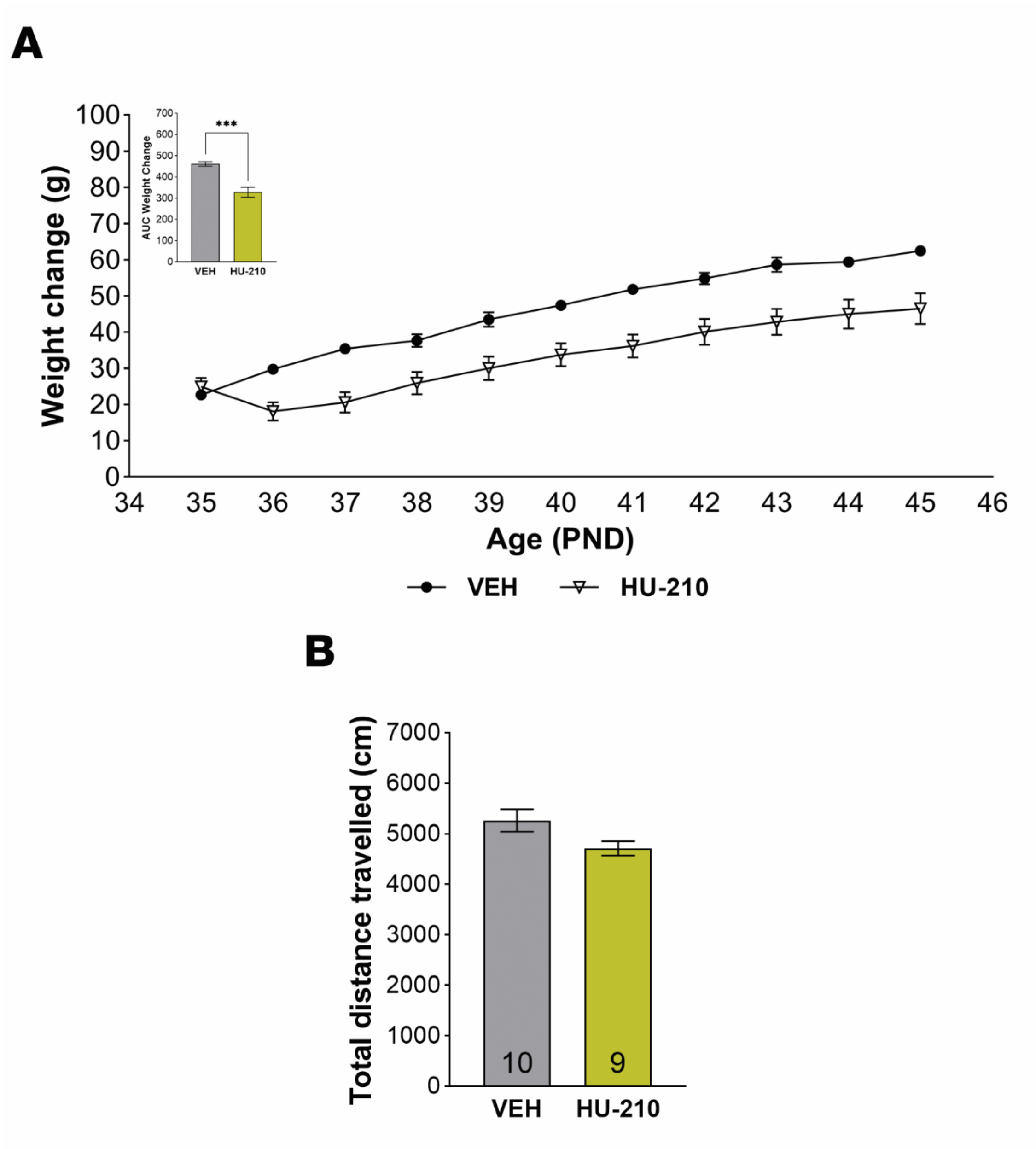


**Supplementary Figure S2. Chronic adolescent HU-210 treatment induces significant decreases in weight-gain, without affecting spontaneous locomotor activity.**

**A.** Changes in animal weight over time. To assess alterations in weight gain, the weight of each animal at PND28 was subtracted to its daily weight during the period of injections (PND35-45). To avoid multiple comparisons, the area under curve was obtained and compared between groups (inset), and evidenced a clear HU-210-induced decrease in weight-gain; **B.** HU-210-treated animals did not significantly differ from vehicle-treated controls with regards to the total distance travelled during either the first or the second OFT trial, suggesting treatment did not induce any lasting changes in locomotor activity. Data are expressed as mean ± SEM, with number of animals *n* indicated at the bottom of plot bars; *** $p \leq 0.001$, unpaired student's t-test. Graphs correspond to the animals in experiment 2 (see text).

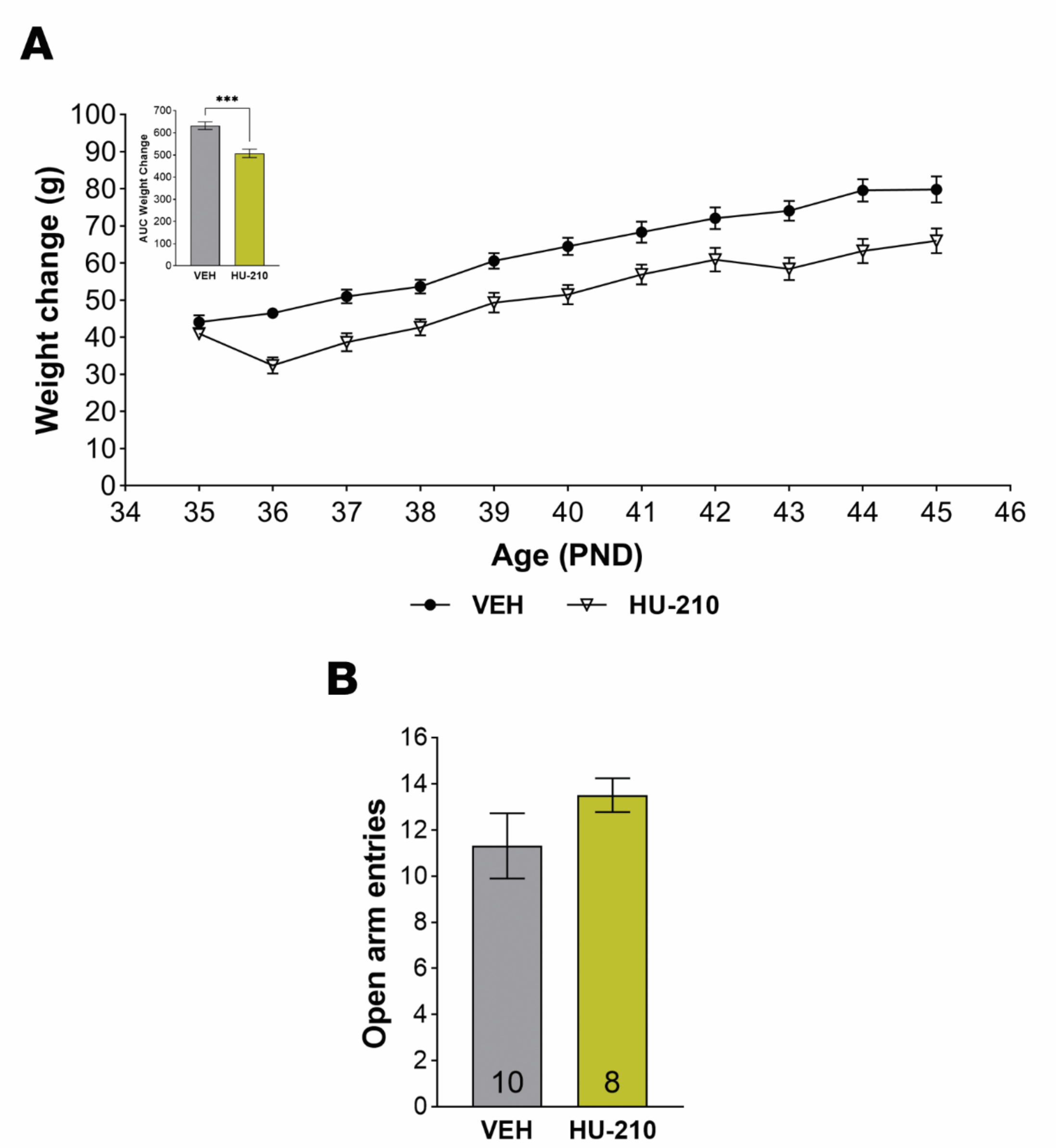


**Supplementary Figure S3. Chronic adolescent HU-210 treatment induces significant decreases in weight-gain, without affecting anxiety-like behavior.**

**A.** Changes in animal weight over time. To assess alterations in weight gain, the weight of each animal at PND28 was subtracted to its daily weight during the period of injections (PND35-45). To avoid multiple comparisons, the area under curve was obtained and compared between groups (inset), and evidenced a clear HU-210-induced decrease in weight-gain; **B.** In accordance with the absence of differences in open arm permanence in the EPM (see fig. 3B), no differences were observed in the number of open arm entries. Data are expressed as mean ± SEM, with number of animals *n* indicated at the bottom of plot bars; *** $p \leq 0.001$, unpaired student's t-test. Graphs correspond to the animals in experiment 3 (see text).

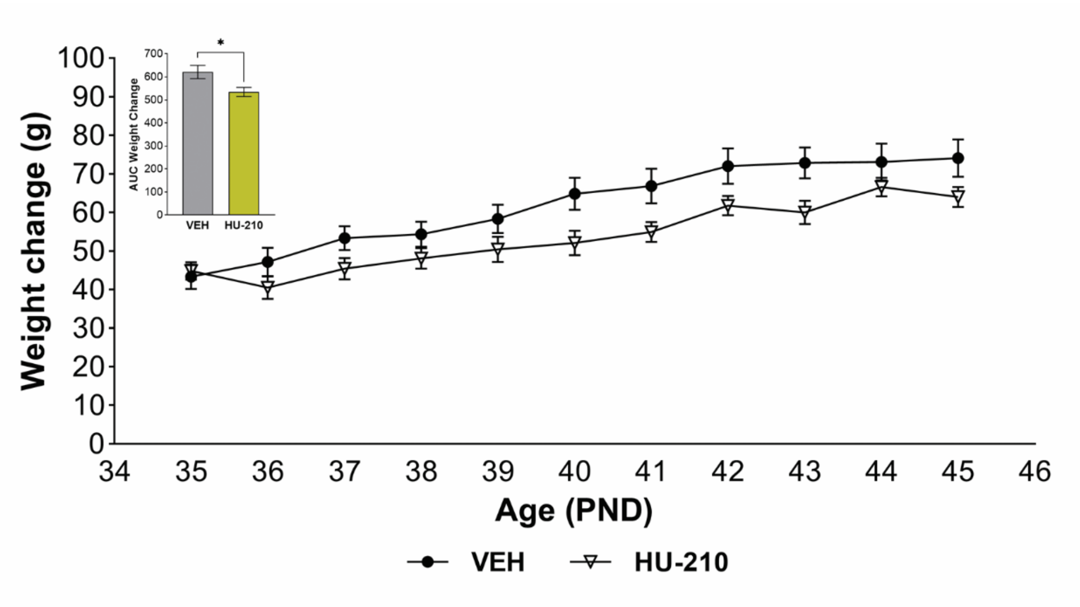


**Supplementary Figure S4. Chronic adolescent HU-210 treatment induces significant decreases in weight-gain.**

Changes in animal weight over time. To assess alterations in weight gain, the weight of each animal at PND28 was subtracted to its daily weight during the period of injections (PND35-45). To avoid multiple comparisons, the area under curve was obtained and compared between groups (inset), and evidenced a clear HU-210-induced decrease in weight-gain. Data are expressed as mean ± SEM; * $p < 0.05$, unpaired student's t-test. Graphs correspond to the animals in experiment 4 (see text).
